# Glucosylceramides impact cellulose deposition and cellulose synthase complex motility in Arabidopsis

**DOI:** 10.1101/2024.03.25.585105

**Authors:** Jose A. Villalobos, Rebecca E. Cahoon, Edgar B. Cahoon, Ian S. Wallace

**Affiliations:** Department of Biochemistry and Molecular Biology, University of Nevada, Reno, Reno, NV 89557; Department of Biochemistry and Molecular Biology, Athens, GA 30602; Complex Carbohydrate Research Center, University of Georgia, Athens, GA 30602; Department of Biochemistry & Center for Plant Science Innovation, University of Nebraska, Lincoln, Lincoln, NE 68588

**Keywords:** Cellulose/ sphingolipids/ chemical biology/ Cellulose Synthase

## Abstract

Cellulose is an abundant component of plant cell wall matrices, and this para-crystalline polysaccharide is synthesized at the plasma membrane by motile Cellulose Synthase Complexes (CSCs). However, the factors that control CSC activity and motility are not fully resolved. In a targeted chemical screen, we identified the alkylated nojirimycin analog *N*-Dodecyl Deoxynojirimycin (ND-DNJ) as a small molecule that severely impacts Arabidopsis seedling growth. Previous work suggests that ND-DNJ-related compounds inhibit the biosynthesis of glucosylceramides (GlcCers), a class of glycosphingolipid associated with plant membranes. Our work uncovered major changes in the sphingolipidome of plants treated with ND-DNJ, including reductions in GlcCer abundance and altered acyl chain length distributions. Crystalline cellulose content was also reduced in ND-DNJ-treated plants as well as plants treated with the known GlcCer biosynthesis inhibitor N-[2-hydroxy-1-(4-morpholinylmethyl)-2-phenyl ethyl]-decanamide (PDMP) or plants containing a genetic disruption in GLUCOSYLCERAMIDE SYNTHASE (GCS), the enzyme responsible for sphingolipid glucosylation that results in GlcCer synthesis. Live-cell imaging revealed that CSC speed distributions were reduced upon treatment with ND-DNJ or PDMP, further suggesting an important relationship between glycosylated sphingolipid composition and CSC motility across the plasma membrane. These results indicated that multiple interventions compromising GlcCer biosynthesis disrupt cellulose deposition and CSC motility, suggesting that GlcCers impact cellulose biosynthesis in plants.

## Introduction

Cellulose is one of the most abundant biopolymers on the planet, and this para-crystalline polysaccharide is a major component of plant cell wall extracellular matrices. Cellulose is synthesized at the plasma membrane of plant cells (Paredez et al., 2006) by Cellulose Synthase Complexes (CSCs), which are comprised of multiple non-redundant Cellulose Synthase A (CESA) subunits that serve as the CSC catalytic subunits (Purushotham et al., 2016; Cho et al., 2017). The CSC also contains accessory subunits, such as the KORRIGAN endoglucanase (Vain et al., 2014), Cellulose Synthase Interactive 1 (Gu et al., 2010; Li et al., 2012), Companion of Cellulose synthase (CC) proteins (Endler et al., 2015), and the glycosylphosphatidylinositol (GPI)-anchored protein COBRA (Borner et al., 2003; Li et al., 2013; Liu et al., 2013). Together, these subunits form a massive molecular machine that extrudes newly synthesized cellulose into the apoplast (Purushotham et al., 2020). Live-cell imaging has revealed that CSC localization is dynamic and complex. Fluorescently labeled CSC subunits reside in the Golgi apparatus, small vesicular compartments known as Microtubule Associated Cellulose Synthase Compartments (MASCs) or Small CESA Compartments (SmaCCs) (Crowell et al., 2009; Gutierrez et al., 2009), and as motile particles at the plasma membrane that move with constant velocities of approximately 250 nm/min (Paredez et al., 2006). Plasma membrane localized puncta are through to represent actively synthesizing CSCs, and the rate of CSC motility has been correlated to cellulose biosynthetic output in numerous inhibitor and genetic studies (Paredez et al., 2006; Debolt et al., 2007; Paredez et al., 2008; Gu et al., 2010). Despite these insights, the mechanistic factors that govern CSC motility and therefore activity at the plasma membrane are not fully elucidated.

Sphingolipids are a unique group of lipids that play major structural roles in plant membrane architecture as well as critical roles in programmed cell death, membrane trafficking, plant pathogen interactions, protein anchoring to the plasma membrane, and developmental processes (Ternes et al., 2011; Gronnier et al., 2016). Sphingolipids consist of a fatty acid (FA) group with an acyl chain ranging from 16 to 26 carbons, an 18-carbon sphingosine base, commonly referred to as a Long Chain Base (LCB) that can be di- (d18) or tri- (t18) hydroxylated, and a structurally variable headgroup. In Arabidopsis, LCBs and FA-CoAs are used as substrates for the ceramide synthase enzymes LONGEVITY ASSURANCE GENE ONE HOMOLOG 1 (LOH1), LOH2, and LOH3 to form ceramides (Markham et al., 2011; Ternes et al., 2011). These enzymes have distinct substrate specificities for different forms of LCBs and FA-CoAs (Kyle et al., 2016). Ceramide FA moieties can be hydroxylated to form hydroxyceramides (hCers), and these sphingolipids often undergo further structural modifications, such as the attachment of glucose at the carbon 1 position of the LCB by GLUCOSYLCERAMIDE SYNTHASE (GCS) (Msanne et al., 2015) to form glucosylceramides (GlcCers). Alternatively, hCers can be used as substrates for the synthesis of Glycosylinositolphosphoceramides (GIPCs) (Rennie et al., 2014; Fang et al., 2016). The elaborate headgroup of GIPCs contains a phosphate attached to the LCB, followed by an inositol residue that is attached by INOSITOL PHOSPHORYLCERAMIDE SYNTHASE (Wang et al., 2008). INOSITOL PHOSPHORYLCERAMIDE GLUCURONOSYLTRANSFERASE 1 (IPUT1) uses UDP-Glucuronic acid as a sugar donor to attach glucuronic acid to this inositol moiety to form the core GIPC structure (Tartaglio et al., 2017), which can be further elaborated by the addition of hexose and pentose sugars (Fang et al., 2016; Ishikawa et al., 2018).

Impaired GIPC biosynthesis causes poor growth and reduced cellulose content in Arabidopsis seedlings (Fang et al., 2016), but the mechanistic basis of this observation is unclear. Based on previous work indicating that sphingolipids participate in membrane trafficking and organization, it is possible that GIPCs or other sphingolipids are required for proper transport of the CSC from the Golgi or vesicles to the plasma membrane. It is also unclear whether synthesis of other sphingolipid structural classes is required for proper cellulose deposition, and this issue is exacerbated by the observation that Arabidopsis mutants which are specifically compromised in glycosylated sphingolipid biosynthesis are gametophytically or seedling lethal (Rennie et al., 2014; Msanne et al., 2015; Tartaglio et al., 2017).

Prior work has highlighted that various molecules based on deoxynojirimycin scaffolds can impact sphingolipid biosynthesis by inhibiting the synthesis or catabolism of these important lipid species (Ashe et al., 2011; Rugen et al., 2018). Here, we assayed a group of nojirimycin analogs against *Arabidopsis thaliana* to identify compounds that caused root growth inhibition and root swelling, which are hallmarks of cell wall biosynthesis inhibition. From this chemical screen, *N-*dodecyl deoxynojirimycin (ND-DNJ) was identified as a robust inhibitor of root growth. Subsequent experiments revealed that ND-DNJ inhibits the biosynthesis of glycosphingolipids, and also inhibits the deposition of crystalline cellulose. These effects were recapitulated in an Arabidopsis mutant that is compromised in GlcCer biosynthesis. Live-cell imaging of CSCs in Arabidopsis seedlings treated with ND-DNJ also revealed that this compound caused reduced CSC motility, suggesting that GlcCers are required for CSC activity and motility.

## Results

### Screening of nojirimycin-based analogs

Prior work has indicated that a select subset of nojirimycin-based compounds significantly impact plant growth and development (Rugen et al., 2018). Based on our prior interest in identifying inhibitors of plant cell wall glycosyltransferases (Xia et al., 2014; Villalobos et al., 2015), we screened a small panel of commercially available nojirimycin analogs (Supplemental Figure 1A) that was largely orthogonal to a previous study (Rugen et al., 2018) for their ability to induce cell wall biosynthetic defects, including reduced root elongation and root swelling. To evaluate the effects of this inhibitor panel, Arabidopsis seedlings were germinated on MS medium containing 100 µM test compound, grown for 7 days as described in Materials and Methods, and their root lengths were quantified (Supplemental Figure 1B). These experiments revealed that *N*-dodecyl-deoxynojirimycin (ND-DNJ) (Figure 1A) robustly inhibited Arabidopsis root growth, while the majority of additional compounds in the screen were only moderately effective or completely inactive in this assay.

**Figure 1:**
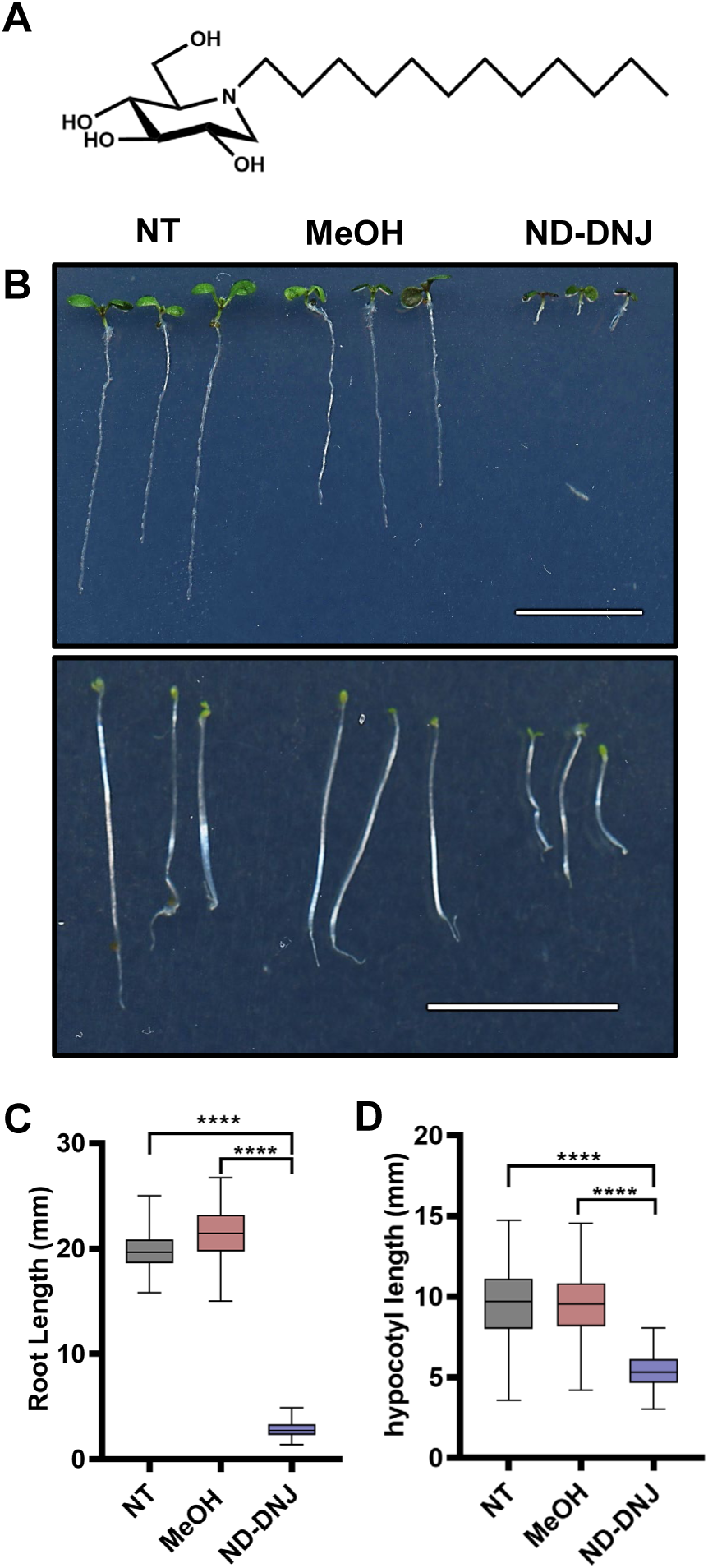
Phenotypic analysis of Arabidopsis seedlings treated with ND-DNJ: (A) The structure of *N*-dodecyl Deoxynojirimycin (ND-DNJ) is shown. (B) Wild-type Col-0 Arabidopsis seedlings were germinated on MS media supplemented with 100 µM ND-DNJ with (top panel) or without (bottom panel) sucrose and grown vertically in the light for 7 days (top panel) or in the dark for 5 days (bottom panel). Scale bars in both panels represent 10 mm. Seven-day-old light-grown root lengths (C) and 5-day old dark-grown hypocotyl lengths (D) of untreated seedlings (NT; gray boxes), seedlings treated with 0.1% methanol (MeOH; red boxes), or 100 µM ND-DNJ (ND-DNJ; blue boxes) were quantified as described in Materials and Methods. In box plots, error bars represent minimum and maximum values, the outer limits of the box represent first and third quartile values, and the center line represents the median value (n = 80-91 in (C) and 100-159 in (D); **** indicates P value < 0.0001 by ANOVA and Tukey post-hoc analysis).

To further investigate the phenotypic defects elicited by ND-DNJ, we performed a series of detailed growth assays. Consistent with the initial screening assays, 100 µM ND-DNJ treatment inhibited Arabidopsis primary root growth by nearly 80% (Figure 1B and 1C) and inhibited Arabidopsis dark-grown hypocotyl elongation by 50% (Figure 1B and 1D). Further detailed microscopy analysis and quantification revealed that 100 µM ND-DNJ treatment caused Arabidopsis primary roots to swell by 30% compared to untreated or solvent controls (Supplemental Figure 2A). A dose-response curve was constructed to measure the effect of increasing ND-DNJ concentrations on primary root elongation (Supplemental Figure 2B). These assays revealed that ND-DNJ inhibited primary root elongation in a dose-dependent manner with an apparent IC_50_ of 21 µM. Overall, these results indicate that ND-DNJ is an effective inhibitor of primary root growth, and potentially suggest that this inhibitor causes defects in cell expansion that are linked to inhibition of cell wall polysaccharide deposition.

### Sphingolipid profiles are altered upon ND-DNJ treatment

Molecules that are structurally similar to ND-DNJ inhibit glucocerebrosidases in animal systems, impact sphingolipid degradation, and serve as an effective drug treatment for Gaucher’s disease as substrate assisted chaperones (Stirnemann et al., 2017). A ceramide analog, N-[2- hydroxy-1-(4-morpholinylmethyl)-2-phenyl ethyl]- decanamide (PDMP) has also been widely used as a GlcCer biosynthesis inhibitor to study vesicular trafficking defects associated with GlcCer depletion (Melser et al., 2010), and a recent screen of nojirimycin-based analogs, which did not include ND-DNJ, also suggested that nojirimycin-based molecules are GlcCer biosynthesis inhibitors (Rugen et al., 2018). Importantly, *N*-5-(adamantane-1-yl-ethoxy) pentyl- L-ido-deoxynojirimycin (L-ido-AEP-DNJ) produced similar phenotypic defects compared to ND-DNJ (Rugen et al., 2018), and we confirmed in our own experiments that PDMP inhibits root and hypocotyl elongation to a similar degree as ND-DNJ (Supplemental Figure 3). Therefore, we postulated that ND-DNJ may inhibit GlcCer biosynthesis in a similar manner to these structurally related molecules.

To test this hypothesis, Arabidopsis seedlings grown for 7 days in the presence of 100 µM ND-DNJ or 50 µM PDMP were lyophilized and subjected to established sphingolipid extraction procedures as described in Materials and Methods. Untreated plants and solvent control seedlings grown in either 0.1% methanol or 0.1% DMSO for ND-DNJ and PDMP, respectively, were processed in the same fashion. The resulting sphingolipid extracts were subjected to targeted lipidomic analysis to quantify the abundance of various sphingolipid metabolites, including free long chain bases (LCBs), GlcCers, GIPCs, hydroxyceramides (hCers), and ceramides (Cers). These experiments revealed that ND-DNJ and PDMP treatment elicited clear impacts on sphingolipid biosynthesis (Figure 2). Arabidopsis seedlings treated with ND-DNJ exhibited a 30% reduction in total GlcCer content compared to untreated controls, with decreases primarily reflected in GlcCers containing d18:1 and d18:2 LCBs (Figure 2D and 2E). In contrast, total GIPC content was minimally impacted by ND-DNJ treatment, with only GIPC species containing d18:1 LCBs exhibiting a significant reduction (Figure 2C and 2E). ND-DNJ treatment also resulted in nearly a 10-fold increase in total free LCB content (Supplemental Figure 4A) with major increases in free d18:0, t18:0, and t18:1 LCBs. Notably, many phosphorylated LCBs were also elevated with d18:1-P, t18:0-P, and t18:1-P being the most prominent (Supplemental Figure 4C). Upon further analysis of d18:1 and t18:1 LCB-containing GlcCers, the most prominent GlcCer species for each LCB, we found that the abundance of d18:1 LCB-containing GlcCers containing C16 acyl groups was reduced by 40% (Supplemental Figure 5) in ND-DNJ treated seedlings. In contrast, t18:1 LCB-containing GlcCers with C16 acyl groups increased in abundance by 50%, suggesting that d18 GlcCers are preferentially impacted by ND-DNJ treatment. We also noted that the concentrations of the highly abundant t18:1 GlcCers with acyl groups ranging from 22 to 26 carbons were minimally impacted by ND-DNJ, but concentrations of d18:1 GlcCers containing longer chain acyl groups were reduced by nearly 5-fold. This analysis supports the hypothesis that ND-DNJ reduced GlcCer content in Arabidopsis.

**Figure 2:**
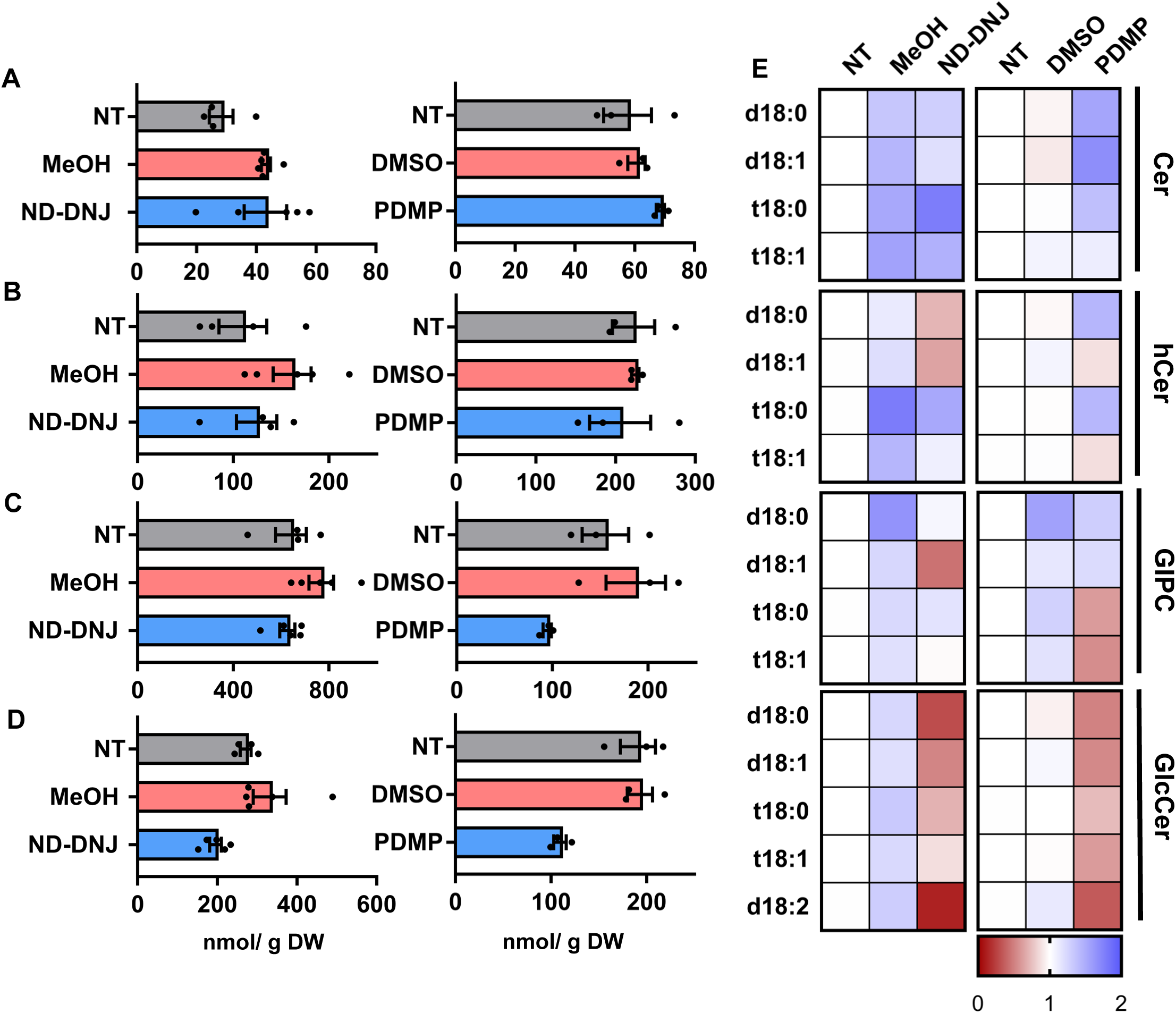
Sphingolipid compositional analysis of Arabidopsis seedlings treated with ND- DNJ and PDMP: Arabidopsis seedlings were propagated on MS media containing either 100 µM ND-DNJ or 50 µM PDMP in the light for 7 days as described in Materials and Methods. MS medium containing either 0.1% methanol (MeOH) or 0.1% DMSO (DMSO) served as solvent controls for ND-DNJ or PDMP treatment, respectively. Each sample set was prepared with independent untreated controls (NT) grown simultaneously with the treatment groups. Sphingolipid species were extracted and quantified as described in Materials and Methods. The total content in nmol/ g DW of (A) ceramides, (B) hydroxceramides, (C) GIPCs, and (D) GlcCers is shown. In all plots, gray bars represent untreated control, red bars represent solvent treatment, and blue bars represent compound treatment. Error bars represent SEM (n = 5 for ND- DNJ treatment set and n = 3 for PDMP treatment set; all values are shown as points). (E). Relative contents of sphingolipid species varying by long-chain base for four major classes of sphingolipids are shown in a heat map representation. The content for each species in untreated controls was set to a value of 1 and treated samples are calculated as a fraction of this value. Warmer colors in the heat map represent relative reductions in specific lipid species, while cooler colors represent relative increases. A scale bar is shown to relate color change to relative quantification.

Consistent with its ascribed function as a GCS inhibitor (Inokuchi et al., 1989; Barbour et al., 1992; Fenderson et al., 1992; Rugen et al., 2018), PDMP treatment caused a 40% reduction in total GlcCer content, with substantial decreases in both d18 (46-65%) and t18 (30-40%) LCB- containing GlcCers (Figure 2D and 2E). PDMP caused only a 1.8-fold increase in total LCB species compared to the 9.3-fold increase elicited by ND-DNJ treatment (Supplemental Figure 4B). Interestingly, GIPC profiles were also impacted by PDMP treatment. Total GIPC content was reduced by 39% in PDMP-treated seedlings, reflected primarily by a reduction in t18:0 and t18:1 LCB species (Figure 2C and 2E). Overall, these results suggest that ND-DNJ and PDMP have complex effects on Arabidopsis sphingolipid content and that the common outcome of treatment with these inhibitors is a substantial reduction in GlcCer species.

### ND-DNJ and PDMP treatments alter cell wall polysaccharide composition profiles

While sphingolipids are required for numerous physiological processes in plants, it is important to note that inhibition of GIPC glycosylation via genetic disruption of Arabidopsis *GIPC MANNOSYL-TRANSFERASE1* caused substantial reduction in crystalline cellulose content (Fang et al., 2016). However, the impact of GlcCer metabolism on cellulose deposition has not been previously examined. Therefore, we analyzed matrix polysaccharide composition and crystalline cellulose content (Villalobos et al., 2015) of 7-day-old seedlings treated with 100 µM ND-DNJ or 50 µM PDMP. Six commonly occurring cell wall monosaccharides were assayed by alditol acetate analysis, and these experiments revealed that arabinose content increased nearly 2- fold in ND-DNJ treated seedlings while the other five cell wall matrix-associated monosaccharides remain relatively unchanged (Figure 3A). A similar elevation of arabinose content was not observed under PDMP treatment (Figure 3B). Intriguingly, total matrix polysaccharide glucose contents of ND-DNJ and PDMP-treated seedlings were nearly five times greater compared to untreated and solvent controls (Figure 3E and 3F). This observation prompted further analysis of the matrix polysaccharide glucose content in ND-DNJ and PDMP treated seedlings. Using 10 mg of the remaining AIR sample, we removed starch using porcine amylase and pullulanase overnight. Enzymes were removed with solubilized glucose, and the remaining tissue was subjected to alditol acetate assay. Glucose content in ND-DNJ and PDMP-treated seedlings was still elevated compared to untreated and solvent controls (Supplemental Figure 6), potentially suggesting that this increase in matrix polysaccharide glucose content is the result of an additional glucan, such as wound-induced callose or non-crystalline cellulose.

**Figure 3:**
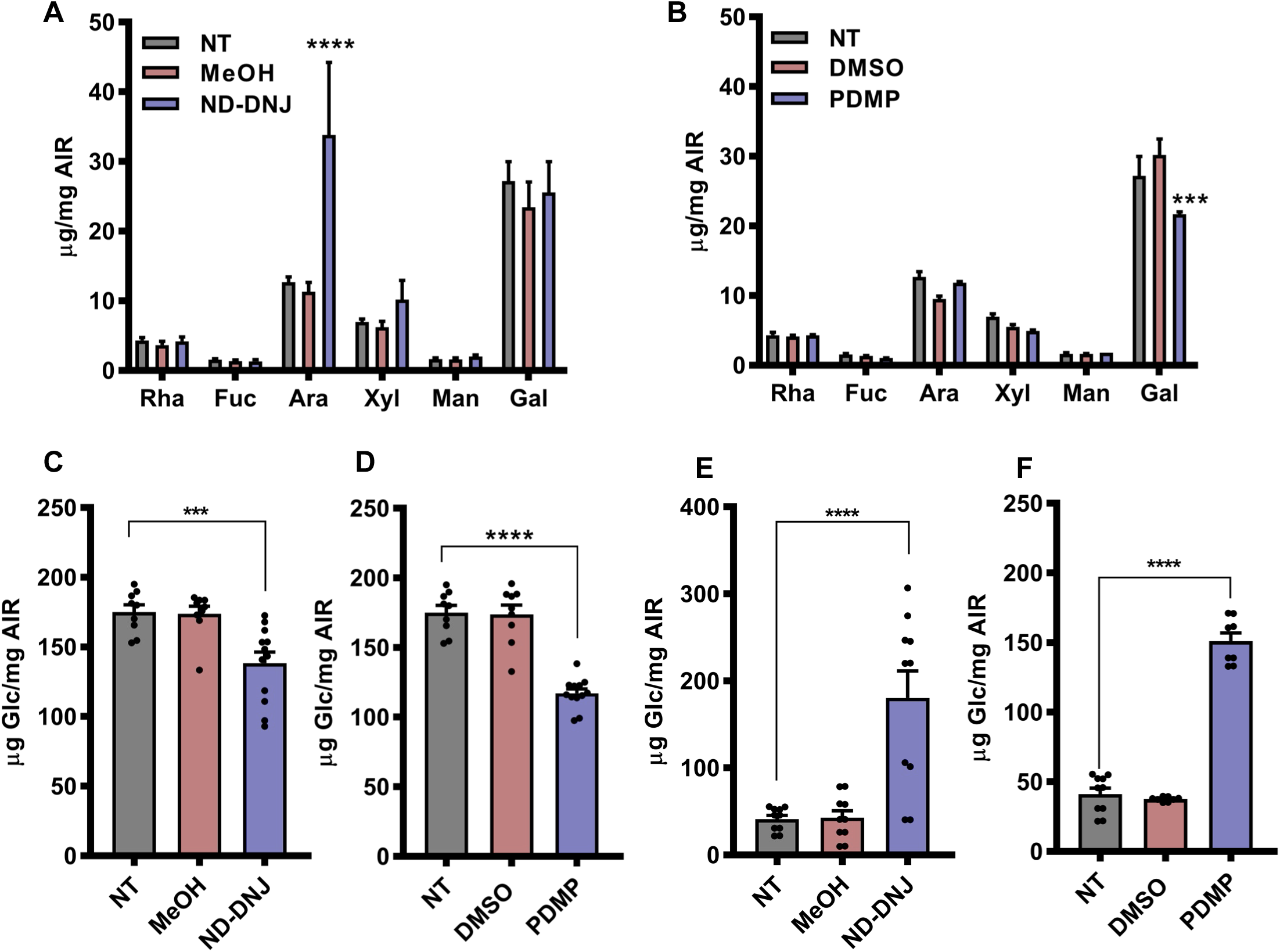
Cell wall compositional changes elicited by ND-DNJ and PDMP treatment: Seven-day-old light-grown Col-0 Arabidopsis seedlings were grown on MS media containing 100 µM ND-DNJ or 50 µM PDMP as indicated. Seedlings grown on MS media alone (NT) or in the presence of 0.1% DMSO (DMSO) or methanol (MeOH) served as negative controls. Alcohol Insoluble Residue (AIR) was prepared from seedlings as described in Materials and Methods. The matrix polysaccharide compositions of seedlings treated with 100 µM ND-DNJ (A) or 50 µM PDMP (B) are shown. Monosaccharides are represented by the following three letter designations: Rha; rhamnose, Fuc; fucose, Ara; arabinose, Xyl; xylose, Man; mannose, Gal; galactose. Crystalline cellulose and matrix polysaccharide glucose contents were also measured for seedlings treated with 100 µM ND-DNJ (C and E) or 50 µM PDMP (D and F). Error bars in each panel represent SEM (n = 8-10). ***, and **** represent P values of < 0.001, and <0.0001 respectively by one-way ANOVA followed by Tukey post-hoc analysis. All data points are shown in C through F.

Crystalline cellulose quantification analysis was also investigated via Updegraff analysis (Updegraff, 1969), and these experiments demonstrated that seedlings germinated under ND- DNJ or PDMP treatment displayed at 25-35% reduction in crystalline cellulose content (Figure 3C and 3D). Overall, these results indicate that ND-DNJ and PDMP treatment cause a significant reduction in crystalline cellulose content with a concomitant increase in matrix polysaccharide glucose content, suggesting that GlcCer biosynthesis is necessary for the proper deposition of crystalline cellulose.

GlcCers are produced via the action of GCS, and only one GCS enzyme has been identified in the Arabidopsis genome (Msanne et al., 2015). *gcs-1* loss-of-function mutants germinate but exhibit seedling growth arrest after several days under light-grown conditions. To investigate whether loss-of-function mutations in the *GCS* gene cause similar effects compared to ND-DNJ or PDMP treatment, we grew the previously described *gcs-1* mutants on MS medium in the light and first measured root elongation. As demonstrated in Figure 4B, *gcs-1* mutant root lengths were reduced by 90% compared to wild-type Col-0 controls. Thus this mutant exhibits similar phenotypic defects compared to ND-DNJ and PDMP treatment.

**Figure 4:**
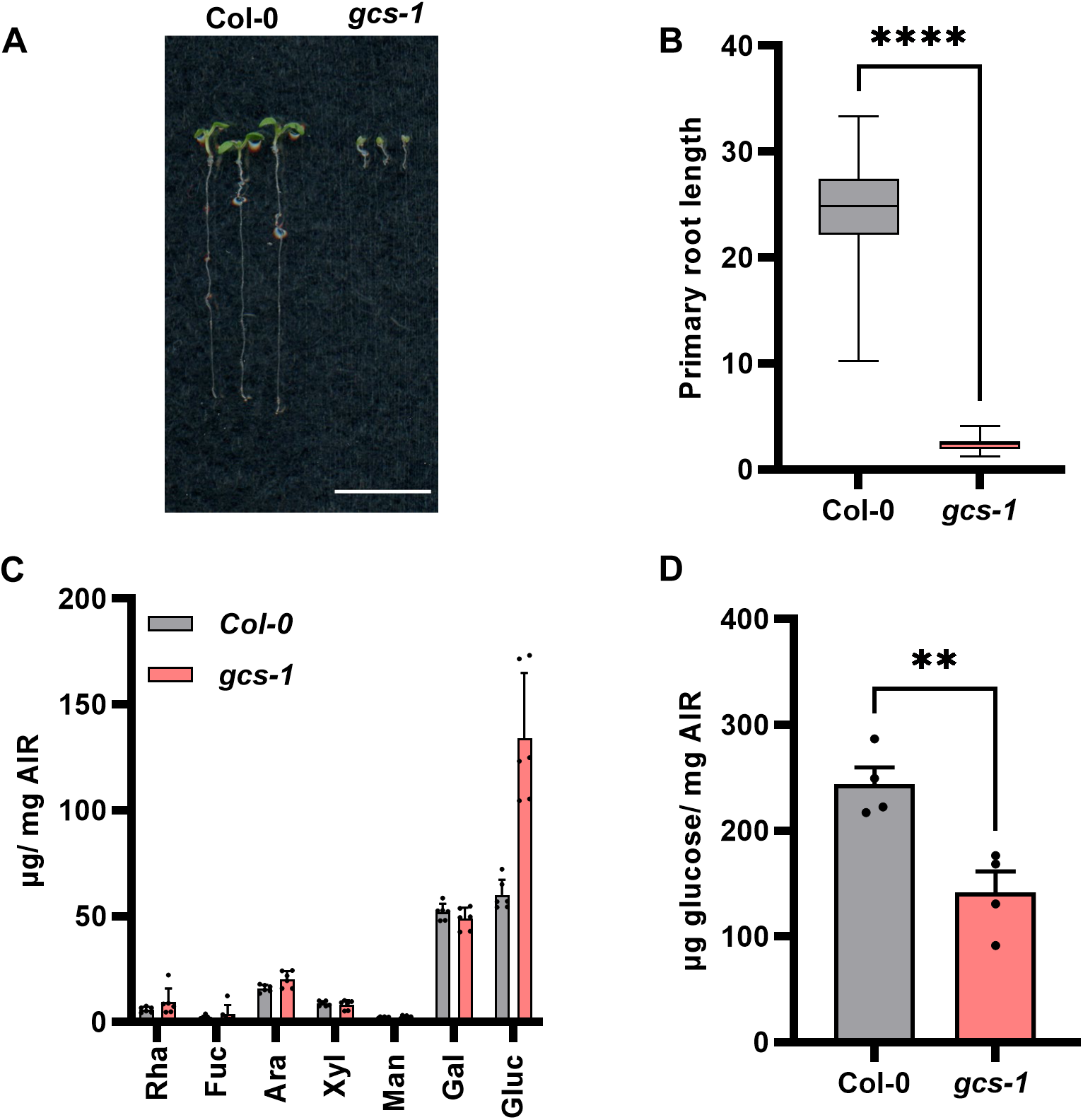
Cell wall compositional analysis of Arabidopsis *gcs* mutant: Arabidopsis Col-0, and *gcs-1* seedlings were grown for 7 days in the light on MS media as described in Materials and Methods. (A) Representative images of 7-day-old Col-0, and *gcs-1* homozygotes are shown. Scale bar represents 10 mm. (B) Quantification of root lengths from 7-day-old light-grown Col-0 (gray box), and *gcs-1* (red box) seedlings. Bars represent minimum and maximum values, the outer limits of the box represent first and third quartile values, and the central line of the box represents the median (n=27-110). Alcohol insoluble residue (AIR) prepared from Col-0 (gray bars), and *gcs-1* (red bars) seedlings as described in Materials and Methods was used to measure the monosaccharide composition of matrix polysaccharides by alditol acetate analysis (C) and crystalline cellulose content (D). For both C and D, error bars represent SEM (n = 4). **, and **** represent P<0.01, and P<0.0001 by Student’s t-test, respectively.

Next, we investigated the cell wall composition of *gcs-1* mutant plants compared to wild-type Col-0 controls. Alcohol Insoluble Residue (AIR) from 7-day-old light grown seedlings was prepared as described in Materials and Methods, and the resulting AIR samples were subjected to alditol acetate monosaccharide analysis as well as crystalline cellulose quantification as described above. These experiments revealed that *gcs-1* mutant plants exhibited 2-fold higher matrix polysaccharide glucose contents, similar to observations of ND-DNJ and PDMP treated plants. The remaining matrix polysaccharide-associated monosaccharides were unchanged in *gcs-1* plants compared to wild-type Col-0 controls (Figure 4C). Crystalline cellulose content of *gcs-1* mutants was also measured (Figure 4D), revealing that the *gcs-1* mutant exhibited a 42% reduction in crystalline cellulose content. Overall, these results suggest that genetic disruption of Arabidopsis *GCS* results in similar phenotypic and cell wall chemistry defects compared to ND- DNJ and PDMP treatment, further supporting the hypothesis that disruption of GlcCer biosynthesis causes the inhibition of cellulose deposition.

### ND-DNJ or PDMP treatment inhibits Cellulose Synthase Complex motility

Cellulose is synthesized by CSCs in the plasma membrane, which can be visualized in living cells by examining the dynamics of fluorescent protein-CSC subunit fusions (Paredez et al., 2006; Debolt et al., 2007; Gutierrez et al., 2009). This imaging system is a useful tool to investigate the mechanism of inhibitors that impact cellulose biosynthesis (Paredez et al., 2006; Debolt et al., 2007; Xia et al., 2014). To examine the impact of PDMP and ND-DNJ treatment on CSCs, we treated dark-grown GFP-CESA3-expressing transgenic seedlings with 100 µM ND- DNJ or 50 µM PDMP for 1 hour and examined CSC speed distributions in hypocotyl epidermal cells of these seedlings via spinning disc confocal microscopy and live-cell cinematography as described in Materials and Methods. Untreated GFP-CESA3 seedlings, or seedlings treated with 0.1% solvent served as controls. As shown in Figure 5A, single-frame images of epidermal cells treated with PDMP or ND-DNJ did not reveal drastic differences in CSC distribution throughout the plasma membrane, suggesting that GlcCer biosynthesis inhibition does not significantly impact the positional distribution or overall number of CSCs at the plasma membrane. However, time-averaged projections of CSC trajectories over a 5-minute time-lapse revealed that CSCs in ND-DNJ and PDMP-treated seedlings showed shorter motility trajectories than corresponding controls. Indeed, Kymograph analysis of individual CSC particles (Figure 5B) revealed that untreated plants exhibited the typical linear constant velocity pattern that is typically observed of motile CSCs, while the CSC of ND-DNJ and PDMP-treated samples exhibited nearly vertical kymographs, suggesting that CSC velocities were reduced under these treatments. We then measured the velocity distribution of CSC particles from 6-10 cells from at least 3 independent seedlings per treatment group (Figure 5C) and found that untreated seedlings exhibited an average speed of 264 nm/min, consistent with previous observations (Paredez et al., 2006; Gutierrez et al., 2009). While this average speed was not significantly impacted by solvent control treatment, average CSC speeds in ND-DNJ and PDMP-treated seedlings were reduced to 107 and 147 nm/min, respectively. We additionally generated multiple *gcs-1* mutant lines expressing fluorescent protein-CESA markers, but unfortunately, *gcs-1* homozygous mutants did not germinate under dark grown conditions, precluding further analysis of CSC dynamics in this genetic background. Overall, these observations indicate that ND-DNJ and PDMP treatment cause decreased CSC motility, suggesting that inhibition of GlcCer biosynthesis by these compounds causes the inhibition of crystalline cellulose deposition by reducing CSC motility and activity.

**Figure 5:**
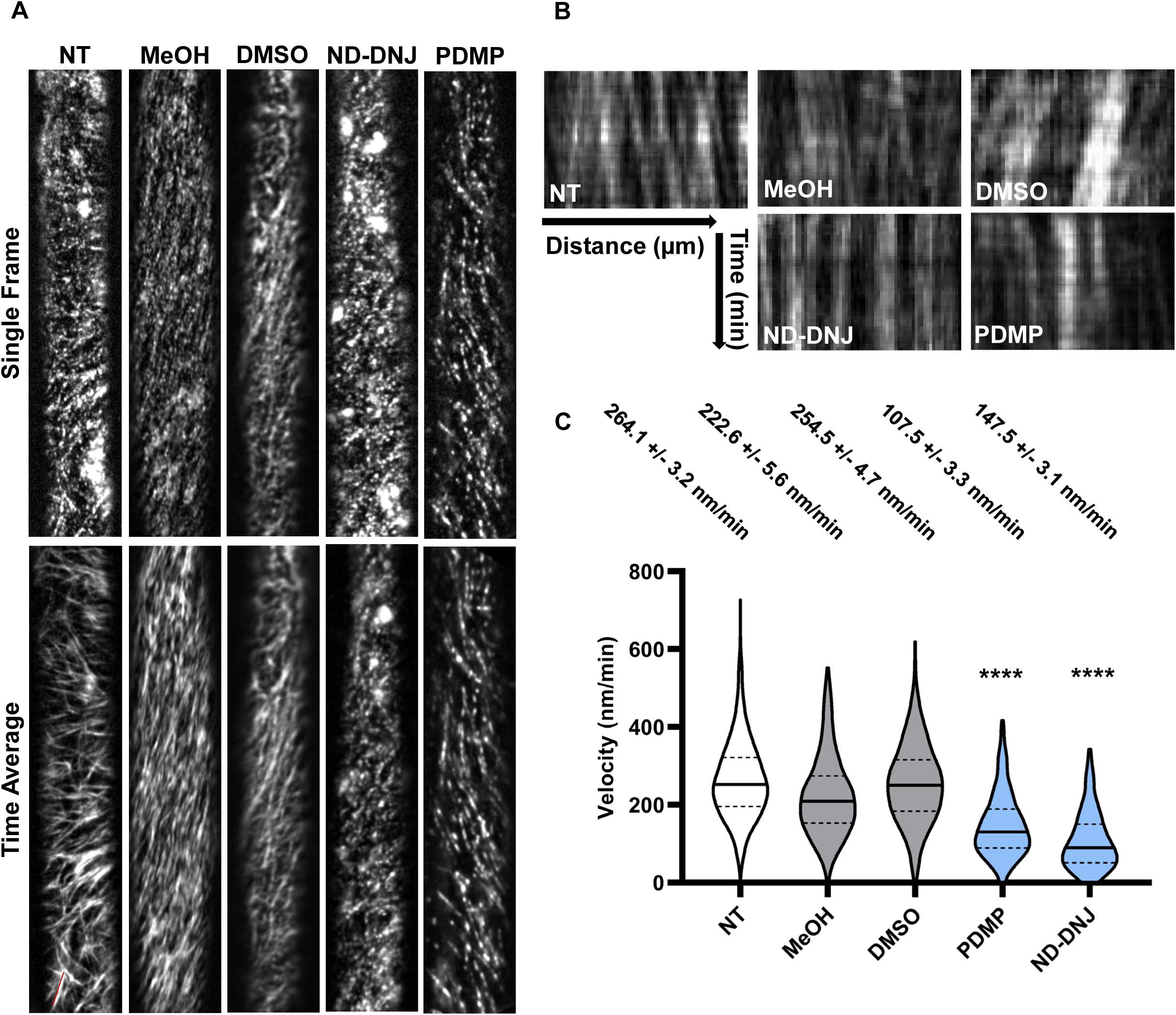
Effects of PDMP and ND-DNJ on the Cellulose Synthase Complex: Transgenic Arabidopsis seedlings expressing pCESA3::GFP-CESA3 were grown in the dark for 4 days as described in Materials and Methods and treated with either 100 µM ND-DNJ or 50 µM PDMP prior to image analysis via spinning disc confocal microscopy. Untreated seedlings (NT) or seedlings treated with 0.1% solvent (DMSO and MeOH) served as controls (A) Representative single frame images (top panels) and corresponding time-average projections of 5-minute videos (bottom panel) are shown for each treatment. (B) Representative kymographs of CESA3 trajectories shown in (A). (C) Quantification of CSC speed distributions. Violin plots showing distribution of CSC velocities around the mean under NT, MeOH, ND-DNJ, DMSO, and PDMP treatments. Average velocities of each treatment are reported above, reported as mean +/- SEM. One-way ANOVA tests were run between each treatment group (**** P<0.0001). N=523-590 velocity measurements across 6-8 cells.

## Discussion

Sphingolipid biosynthesis impacts numerous aspects of plant growth and development, and previous work has suggested a functional relationship between the synthesis of GIPCs and cellulose deposition (Fang et al., 2016), but the mechanistic details of this relationship are unclear and the impact of other sphingolipids on cellulose deposition has not been investigated. Here, we demonstrate that ND-DNJ in an inhibitor of glycosylated sphingolipid biosynthesis (Figure 2), which primarily impacts synthesis of GlcCers and LCBs. Treatment of Arabidopsis seedlings with this compound caused significant phenotypic abnormalities that are commonly associated with cell wall biosynthesis inhibition (Figure 1; Supplemental Figure 2). Treatment of Arabidopsis seedlings with ND-DNJ or the previously described GlcCer biosynthesis inhibitor PDMP (Inokuchi et al., 1989; Barbour et al., 1992; Fenderson et al., 1992; Rugen et al., 2018) also caused defects in cell wall polysaccharide deposition, including reduced crystalline cellulose content and increased cell wall matrix glucose content (Figure 3). Genetic disruption of *GCS* recapitulated these biochemical defects (Figure 4), strongly suggesting that GlcCers are required for normal crystalline cellulose deposition in Arabidopsis. Finally, the mechanism of cellulose biosynthesis inhibition elicited by ND-DNJ and PDMP treatment was investigated by examining the dynamics of CSCs via live-cell imaging, and these experiments revealed that CSC motility, and likely activity, was impaired upon treatment with ND-DNJ or PDMP (Figure 5). Overall, these results suggest that reduced GlcCer content negatively impacts CSC motility and the deposition of crystalline cellulose in the plant cell walls of Arabidopsis.

Both ND-DNJ and PDMP have substantial impacts on GlcCer metabolism, but differences between the detailed sphingolipid species profiles in plants treated with these inhibitors indicate that these molecules may have unique mechanisms of action. PDMP is a well-established inhibitor of GlcCer biosynthesis in metazoan systems (Inokuchi et al., 1989; Barbour et al., 1992; Fenderson et al., 1992), and this compound has been used in plant systems to investigate the impact of GlcCer depletion on various cellular and physiological processes (Melser et al., 2010; Krüger et al., 2013; Rugen et al., 2018). Here, we have demonstrated that PDMP reduces total GlcCer content and impacts both d18 and t18 LCB-containing species to a similar degree, consistent with the idea that this compound inhibits GCS in Arabidopsis. The majority of other sphingolipid species were not substantially impacted by PDMP treatment.

While ND-DNJ elicited similar effects on GlcCer contet, it also caused a nearly 10-fold increase in total free LCB content, and the majority of these more abundant species are t18 LCB-related metabolites. This effect was not observed upon PDMP treatment, suggesting that ND-DNJ and PDMP both reduce total GlcCer content, but potentially through different mechanisms.

We used chemical genetic approaches and affinity purification techniques to attempt the identification of the ND-DNJ target, but despite our best efforts, these approaches were not successful. Thus, the molecular target of ND-DNJ remains to be identified, which is a limitation of this study and will be the focus of future work. However, we suggest that the observed sphingolipid metabolism defects could be explained by ND-DNJ activating the de-acylation or degradation of GlcCers. In this model, GlcCer specific ceramidases or de-glucosylation of GlcCers would occur through the action of glucosylceramidases. The first of these activities has not been described to our knowledge in plants, but a small family of four glucosylceramidase enzymes were recently described in the Arabidopsis genome (Dai et al., 2020). Recombinant Arabidopsis GLUCOSYLCERAMIDASE3 (GCD3) preferentially hydrolyzed glucose moieties of d18 and t18 GlcCers with acyl chains between 18-20 carbons (Dai et al., 2020), which could explain the preferential inhibition of d18 LCB-containing GlcCer synthesis upon ND-DNJ treatment. Loss-of-function mutants in *GCD3* did not exhibit observable phenotypes and did not recapitulate the phenotypes exhibited upon ND-DNJ treatment, but it is important to note that multiple isoforms of GCD were identified, and these genes could be partially redundant or may need to be overexpressed to phenocopy the impacts of ND-DNJ. Importantly, Arabidopsis GCD enzymes are the plant homologs of mammalian glucocerebrosidases, and these enzymes are known to be allosterically activated (or inhibited) by a broad range of imino sugar-based small molecules (Kopytova et al., 2021), suggesting that GCD allosteric activation by ND-DNJ is a plausible mechanism to explain the associated impacts of this molecule. While the mechanism of ND-DNJ remains unknown and will be the subject of future investigations, it is important to note that PDMP treatment and the *gcs-1* mutant exhibited reduced crystalline cellulose content and PDMP elicits similar CSC motility defects compared to ND-DNJ, thus supporting the hypothesis that GlcCer content is important for cellulose deposition.

The inhibition of GlcCer metabolism strikingly reduced speed distribution of CSCs treated with ND-DNJ or PDMP, suggesting that GlcCer content impacts CSC motility and activity. We postulate that this effect could arise from one of at least two possible models. First, GlcCers and other sphingolipids are enriched in membrane nanodomains within the plasma membrane that scaffold protein complexes, and prior work has demonstrated that d18 LCB- containing sphingolipids are particularly enriched in these membrane nanodomains (Borner et al., 2005; Carmona-Salazar et al., 2021) along with cellulose synthase activity (Bessueille et al., 2009). Therefore, a plausible explanation for the reduced speed distribution of CSCs upon GlcCer depletion is that the CSC is encased in a lipid nanodomain of sphingolipids within the plasma membrane, and that disruption of this nanodomain causes the reduced velocity of the CSC. GPI-anchored proteins are often preferentially enriched in membrane nanodomains (Borner et al., 2003; Borner et al., 2005), and the GPI-anchored protein COBRA (COB) is genetically required for cellulose biosynthesis (Roudier et al., 2005). Previous work suggest that COB is specifically required for the deposition of crystalline cellulose (Liu et al., 2013), suggesting that the disruption of GlcCer metabolism could disassociate COB from the CSC and cause the CSC to deposit cellulose with reduced crystallinity. Overall, the results presented here indicate that reducing Arabidopsis GlcCer content via chemical and genetic approaches compromises CSC activity, motility, and the deposition of crystalline cellulose, thus suggesting that these sphingolipids may actuate aspects of plant growth and development by regulating cellulose deposition.

## Materials and Methods

### General plant growth, maintenance, and drug treatments

*Arabidopsis thaliana* Columbia (Col-0) seeds were sterilized in seed sterilization solution (30% [v/v] bleach, 0.1% SDS [w/v]) for 20 minutes at 25 ℃. This solution was removed, and seeds were washed 5 times in sterile water followed by incubation at 4 °C for 48 hours. Seeds were germinated on MS media (1/2 X Murashige and Skoog salts, 10 mM MES-KOH pH 5.7, 1% Sucrose [w/v], and 1% phytoagar [w/v]). *N*-Dodecyl-Deoxynojirymycin (ND-DNJ) and all other compounds described in this study were purchased from Toronto Research Chemicals (TRC) (Ontario, Canada) and added to MS agar media at specified concentrations after the media was autoclaved and cooled to below 50° C. Seedlings were grown under long day conditions (16hr light, 8hr dark) at 22 ℃ vertically for 7 days. Seedling roots were straightened and scanned with a benchtop scanner, root lengths were quantified using ImageJ. For assays involving dark-grown seedlings, MS media lacking sucrose was supplemented with ND-DNJ or PDMP at the indicated concentrations. Seeds were sterilized as described above, plated onto MS media lacking sucrose and incubated in the light for 1 hour. Plates were subsequently wrapped in aluminum foil and grown under long day conditions at 22 °C as described above for 5 days. Etiolated hypocotyl lengths were quantified using ImageJ.

The *gcs-1* (SK-2634) mutant allele was obtained from the Arabidopsis Biological Resource Center. Genomic DNA extraction and PCR genotyping was performed as previously described (Smith et al., 2018). The *gcs-1* mutant allele genotyping was conducted with the following primer set: SK-2634 LP: CCTCATCCTTCGGTCAATATG, SK-2634 RP: GGGACTCTTGGGAGTTATTGC, and SK T-DNA LB: ATACGACGGATCGTAATTTGTCG.

### Analysis of plant cell wall polysaccharides

Alcohol Insoluble Residue (AIR) was prepared from 7-day-old light-grown Arabidopsis seedlings, and this material was used to perform alditol acetate analysis of cell wall matrix polysaccharides as previously described (Villalobos et al., 2015). Crystalline cellulose content was quantified by the Updegraff assay (Updegraff, 1969) with modifications. Five hundred microliters of Updegraff reagent ([8:1:2] [v:v:v] Acetic acid: Nitric acid: water) was added to 10 mg of AIR material, and the samples were heated at 100 ℃ for 30 minutes. Samples were centrifuged at 14,000 x g for 15 minutes at 25 °C, and the supernatant was removed. The pellet was washed with 750 µL of water, followed by centrifugation under the same conditions. The supernatant was removed, and the resulting pellet was air dried at 80 °C. Samples were resuspended in 175 µL of 72% [v/v] H_2_SO_4_ and incubated at 22 °C for 30 minutes. Samples were vortexed and incubated at 22 °C for an additional 15 minutes, followed by dilution with 825 µL of water, and 200 µL of each sample was transferred to a 1.5 mL tube. Four hundred microliters of freshly prepared Anthrone reagent (2% [w/v] Anthrone in concentrated H_2_SO_4_) was added to each sample and heated to 80 ℃ for 10 minutes, then cooled to 22 ℃. Seventy-five microliters of each sample was arrayed into a 96-well flat bottom plate, and sample absorbance was measured at 625 nm in a SpectraMax M5 spectrophotometer. Glucose content in each sample was calculated based on linear regression from glucose standards.

### Sphingolipid quantification

*Arabidopsis* seedlings were germinated in 1/2 X MS + 1% Sucrose media without supplementation or containing 0.1% Methanol, 100 µM ND-DNJ, 0.1% DMSO, or 50 µM PDMP. Seedlings were grown for 7 days under long day conditions in the light as described above, and then transferred to a 15 mL tube, flash frozen in liquid nitrogen, and lyophilized using a Labconco freeze dry system. Sphingolipid extraction and targeted lipidomic analysis were performed using LC-MS/MS as previously described (Markham and Jaworski, 2007; Kimberlin et al., 2013)

### Cellulose synthase complex imaging

A transgenic Arabidopsis line expressing GFP-CESA3 under its native promoter was previously described (Gutierrez et al., 2009). This line was grown on MS agar without sucrose for 4 days in the dark, and etiolated seedlings were mounted in 200 µL of water, 0.1% MeOH, 100 µM ND-DNJ, 0.1% DMSO, or 50 µM PDMP for 1 hour prior to imaging. Seedlings were mounted as previously described (Paredez *et al*., 2006) and imaged on a Leica DMI-8 confocal microscope featuring a CSU-W1 spinning disk head (Yokogawa), a 100x 1.47NA oil immersion objective (Leica), and an iXon Life EMCCD camera (Andor Technology) controlled by VisiView software (Visitron Systems). GFP fluorescence was excited with a 488 nm laser and emission was recorded with a 525 nm filter. CSC motility assays were performed by recording 5-minute time-lapse movies in 10 second increments.

All time-lapse recordings were processed using identical processing conditions in Fiji (https://imagej.net/Fiji). Movies were corrected for drift using rigid body alignment with the StackReg plugin (Thevenaz et al., 1998). Time-lapse images were further processed by performing a 4-frame walking average, background subtraction of 50 pixels, and then contrast enhancement. Velocities were analyzed using the “kymograph evaluation” plug-in of Fiji as previously described (Verbančič et al., 2021). Approximately 100 velocity measurements were collected from each cell, and cells from at least 6-10 seedlings were used to produce CSC speed distributions.

## Author contributions

ISW, JAV, and REC conceived the original research. ISW, JAV, and REC performed the experiments. JAV, ISW, and EBC wrote the manuscript.

---

**Supplemental Figure 1:**
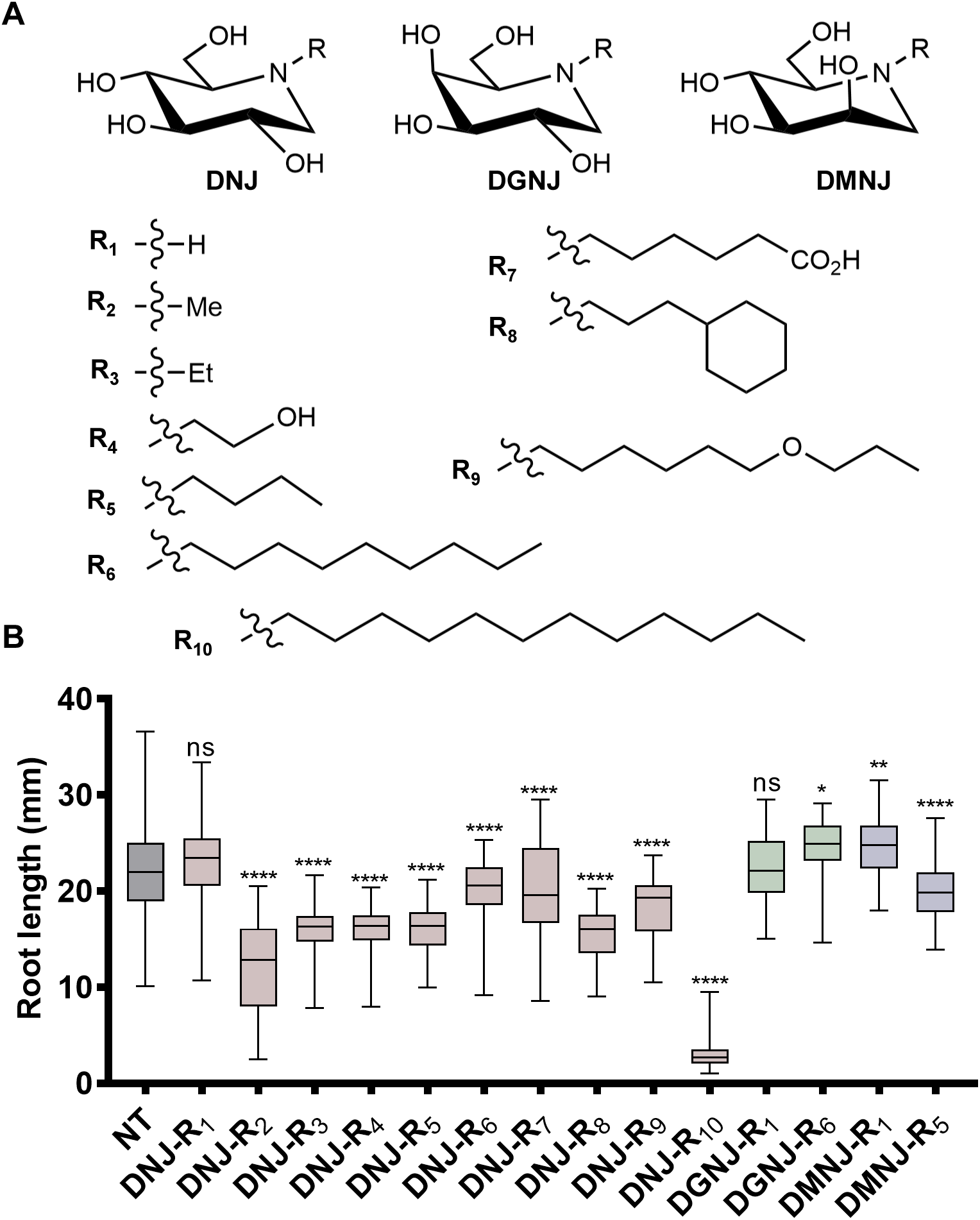
Screen of deoxynojirimycin analogs for impacts on Arabidopsis root growth: (A) The structures of the parent sugar analogs deoxynojirimycin (DNJ), deoxygalactonojirimycin (DGNJ), and deoxymannonojirimycin (DMNJ) are shown as well as the structures of variable R groups attached to the ring nitrogen that were used in this study. (B) Arabidopsis Col-0 seedlings were grown on MS media supplemented with 100 µM test compound for 7 days in the light, and root lengths were measured as described in Materials and Methods. Untreated seedlings (NT) served as controls. X-axis labels indicate the combination of parent sugar analog and R group. DNJ-based analogs are colored in red, DGNJ in green, and DMNJ in blue. Here, N-dodecyldeoxynojirimycin (ND-DNJ) is represented as DNJ-R_10_. Boxes represent first and third quartile, the center lines indicate median values, and bars represent minimum and maximum values (n = 38-109). *, **, and **** indicate statistical difference compared to non-treated controls (P <0.05, P < 0.01, and P < 0.0001 respectively; ns = not significant) by one-way ANOVA followed by Dunnett’s Multiple Comparison analysis.

**Supplemental Figure 2:**
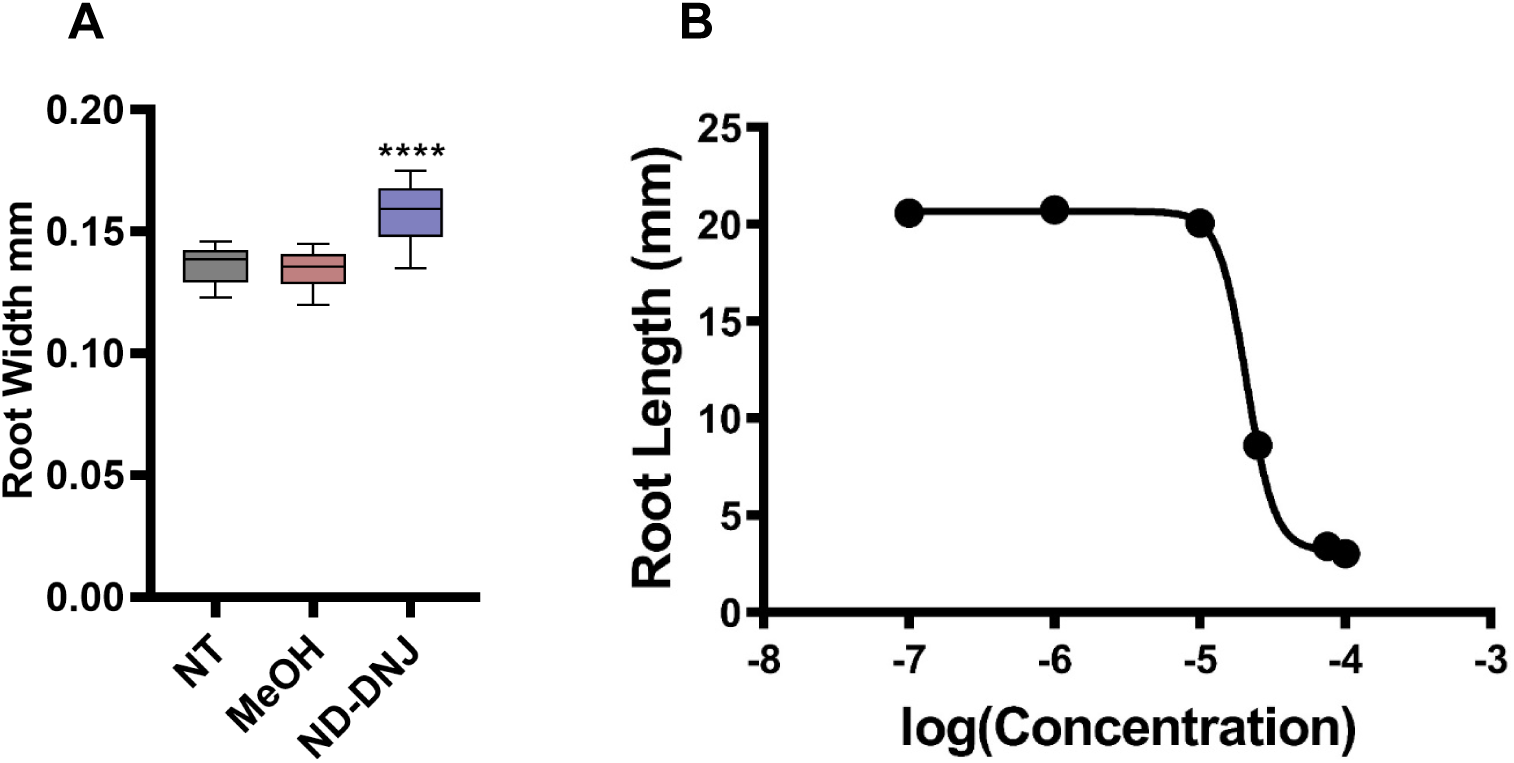
Additional phenotypic effects of ND-DNJ on Arabidopsis seedlings: (A) Arabidopsis seedlings were grown in the light for 7 days on normal MS media (NT; gray boxes), MS media containing 0.1% methanol (MeOH; red boxes), or MS media supplemented with 100 µM ND-DNJ (blue boxes). Seedlings were imaged by confocal microscopy under a 20X objective, and the root width was measured 0.5 mm from the root tip in ImageJ. Boxes represent first and third quartile, the center lines indicate median values, and bars represent minimum and maximum values (n = 12; **** indicates statistical significance (P<0.0001) by One-way ANOVA and Tukey’s post-hoc analysis). (B) Root lengths of 7-day-old light grown seedlings grown on MS media supplemented with increasing concentrations of ND-DNJ were measured in ImageJ as described in Materials and Methods to generate a dose-response curve for ND-DNJ. Error bars represent SEM (n = 44-70).

**Supplemental Figure 3:**
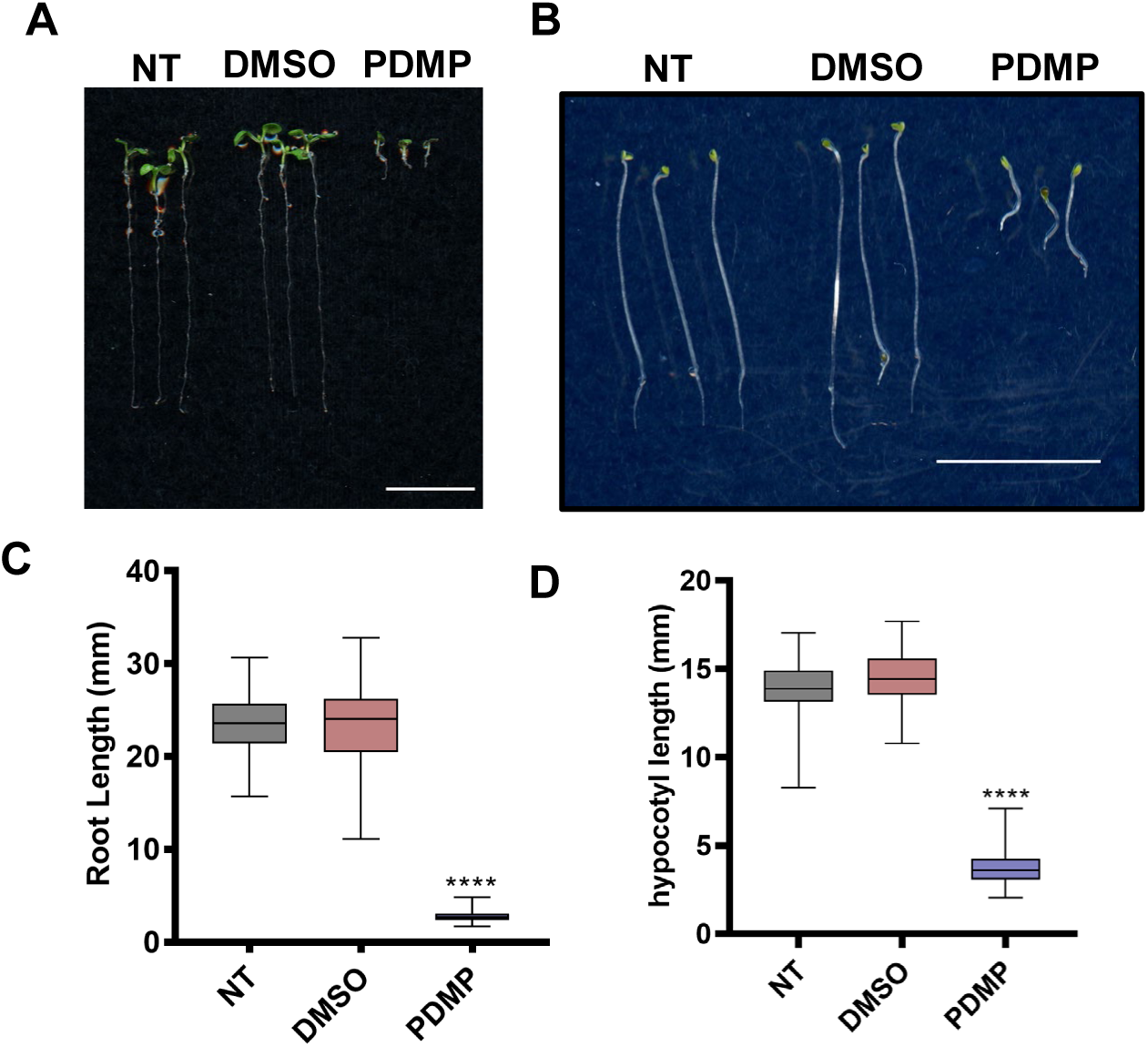
Phenotypic analysis of Arabidopsis seedlings treated with PDMP: Arabidopsis seedlings were grown for 7 days in the light (A) or 5 days in the dark (B) on normal MS media (NT), MS media containing 0.1% DMSO (DMSO) or MS media supplemented with 50 µM PDMP. Scale bar in panels A and B represents 10 mm. Primary root lengths of 7-day-old light grown seedlings (C) and hypocotyl lengths of 5-day-old dark grown Col-0 seedlings (D) grown on MS medium (NT; gray boxes), MS media containing 0.1% DMSO (DMSO; red boxes), and MS media supplemented with 50 µM PDMP (PDMP; blue boxes) were measured as described in Materials and Methods. Boxes represent first and third quartile, the center lines indicate median values, and bars represent minimum and maximum values (n = 121-139 for C and 74-88 for D; **, ***, and **** represent P values of < 0.01, < 0.001, and <0.0001 respectively by one-way ANOVA followed by Tukey post-hoc analysis).

**Supplemental Figure 4:**
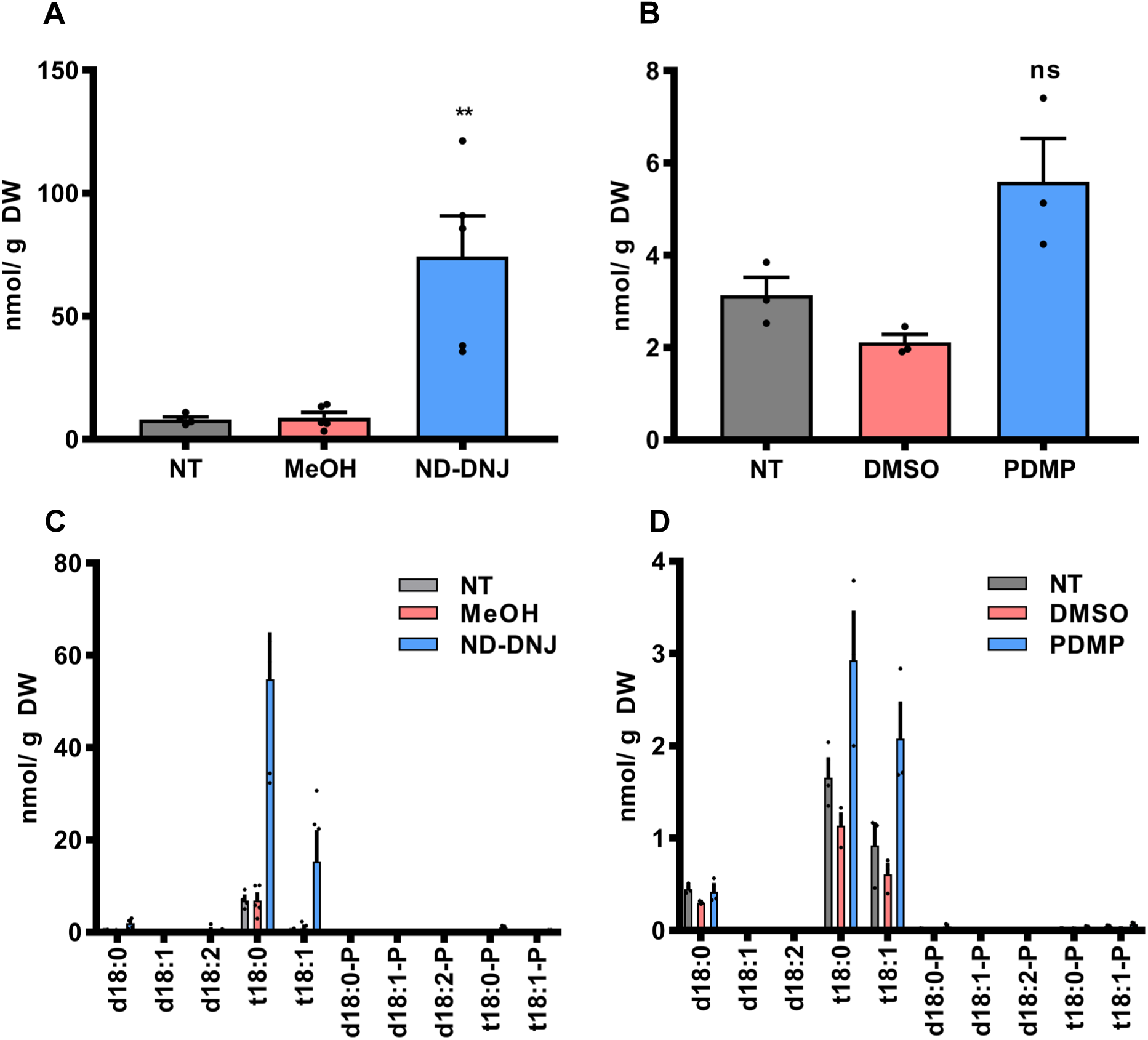
LCB profiles of ND-DNJ and PDMP treated seedlings: Sphingolipids were extracted from 7-day-old Arabidopsis seedlings grown in the light on normal MS media (gray bars), MS media supplemented with 0.1% solvent control (red bars), or MS media containing 50 µM PDMP or 100 µM ND-DNJ (blue bars). Total LCB contents for ND- DNJ-treated (A) and PDMP-treated (B) seedlings are shown as well as the content of specific LCB species in ND-DNJ-treated (C) and PDMP-treated (D) seedlings. Error bars represent SEM (n=3-5). ** indicates P<0.001 by one way ANOVA and Tukey’s post-hoc analysis; ns = not significant. All data points are shown.

**Supplemental Figure 5:**
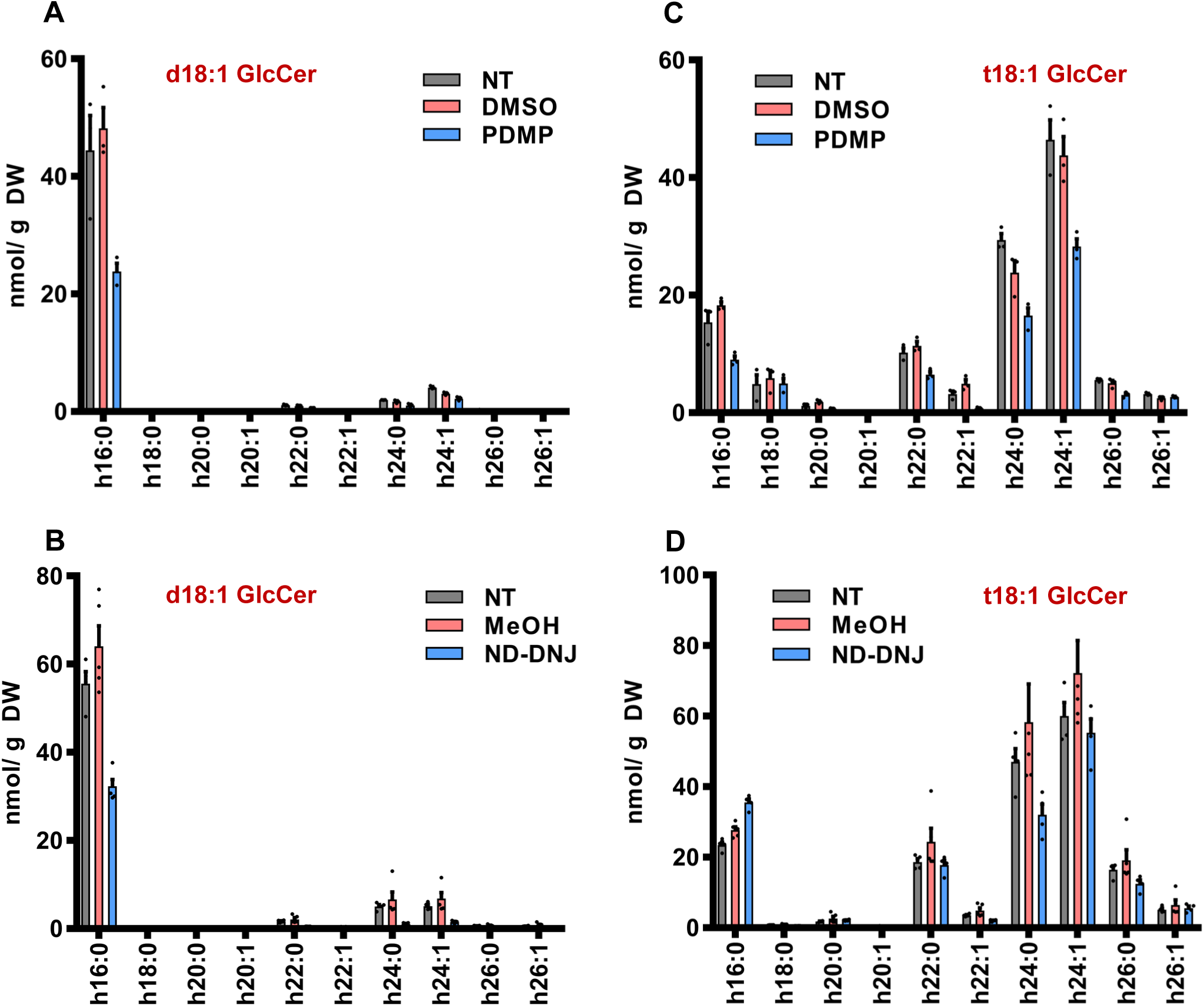
Sphingolipid profiles of select GlcCer species from ND-DNJ and PDMP-treated plants: Sphingolipids were extracted from 7-day-old Arabidopsis seedlings grown in the light on normal MS media (gray bars), MS media supplemented with 0.1% solvent control (red bars), or MS media containing 50 µM PDMP or 100 µM ND-DNJ (blue bars). The acyl group profiles of d18:1 LCB-containing GlcCers from PDMP-treated (A) and ND-DNJ- treated (B) seedlings are shown. Similar profiles of acyl groups from t18:1 LCB-containing GlcCers for PDMP-treated (C) and ND-DNJ-treated (D) seedlings are also shown for comparison. Error bars represent SEM (n = 3-5). All data points are shown.

**Supplemental Figure 6:**
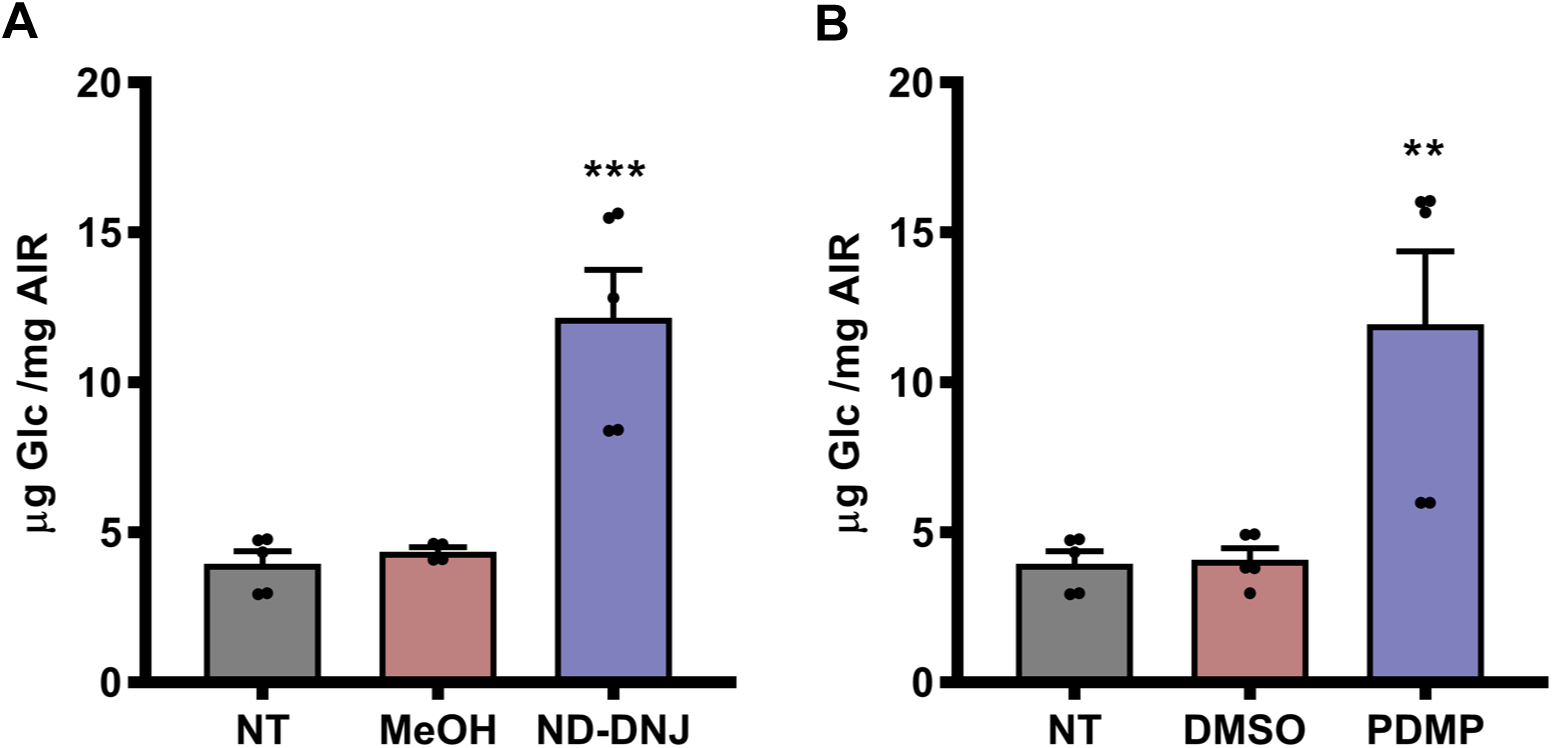
Matrix polysaccharide glucose content after de-starching treatment: Col-0 seedlings were grown in the light for 7 days on MS media containing (A) 100 µM ND-DNJ or (B) 50 µM PDMP. Untreated seedlings (gray bars), or solvent control seedlings (red bars) treated with 0.1% MeOH (A) or 0.1% DMSO (B) served as negative controls. Alcohol Insoluble Residue (AIR) prepared from these seedlings was subjected to enzymatic de-starching, polysaccharide hydrolysis, and alditol acetate analysis as described in Materials and Methods. The matrix polysaccharide glucose contents after de-starching are shown. Error bars represent SEM (n = 5). **, and *** represent P values of < 0.01, and < 0.001 respectively by one-way ANOVA followed by Tukey post-hoc analysis. All data points are shown.

## References

Ashe KM, Bangari D, Li L, Cabrera-Salazar MA, Bercury SD, Nietupski JB, Cooper CGF, Aerts JMFG, Lee ER, Copeland DP, Cheng SH, Scheule RK, Marshall J (2011) Iminosugar-Based Inhibitors of Glucosylceramide Synthase Increase Brain Glycosphingolipids and Survival in a Mouse Model of Sandhoff Disease. PLoS ONE 6: e21758

Barbour S, Edidin M, Felding-Habermann B, Taylor-Norton J, Radin NS, Fenderson BA (1992) Glycolipid depletion using a ceramide analogue (PDMP) alters growth, adhesion, and membrane lipid organization in human A431 cells. Journal of Cellular Physiology 150: 610–619

Bessueille L, Sindt N, Guichardant M, Djerbi S, Teeri TT, Bulone V (2009) Plasma membrane microdomains from hybrid aspen cells are involved in cell wall polysaccharide biosynthesis. Biochemical Journal 420: 93–103

Borner GHH, Lilley KS, Stevens TJ, Dupree P (2003) Identification of Glycosylphosphatidylinositol-Anchored Proteins in Arabidopsis. A Proteomic and Genomic Analysis. Plant Physiology 132: 568–577

Borner GHH, Sherrier DJ, Weimar T, Michaelson LV, Hawkins ND, Macaskill A, Napier JA, Beale MH, Lilley KS, Dupree P (2005) Analysis of Detergent-Resistant Membranes in Arabidopsis. Evidence for Plasma Membrane Lipid Rafts. Plant Physiology 137: 104–116

Carmona-Salazar L, Cahoon RE, Gasca-Pineda J, González-Solís A, Vera-Estrella R, Treviño V, Cahoon EB, Gavilanes-Ruiz M (2021) Plasma and vacuolar membrane sphingolipidomes: composition and insights on the role of main molecular species. Plant Physiology 186: 624–639

Cho SH, Purushotham P, Fang C, Maranas C, Díaz-Moreno SM, Bulone V, Zimmer J, Kumar M, Nixon BT (2017) Synthesis and Self-Assembly of Cellulose Microfibrils from Reconstituted Cellulose Synthase. Plant Physiology 175: 146–156

Crowell EF, Bischoff V, Desprez T, Rolland AL, Stierhof Y-D, Schumacher K, Gonneau M, HöFte H, Vernhettes S (2009) Pausing of Golgi Bodies on Microtubules Regulates Secretion of Cellulose Synthase Complexes in Arabidopsis. The Plant Cell 21: 1141–1154

Dai G-Y, Yin J, Li K-E, Chen D-K, Liu Z, Bi F-C, Rong C, Yao N (2020) The Arabidopsis AtGCD3 protein is a glucosylceramidase that preferentially hydrolyzes long-acyl-chain glucosylceramides. Journal of Biological Chemistry 295: 717–728

Debolt S, Gutierrez R, Ehrhardt DW, Somerville C (2007) Nonmotile Cellulose Synthase Subunits Repeatedly Accumulate within Localized Regions at the Plasma Membrane in Arabidopsis Hypocotyl Cells following 2,6-Dichlorobenzonitrile Treatment. Plant Physiology 145: 334–338

Endler A, Kesten C, Schneider R, Zhang Y, Ivakov A, Froehlich A, Funke N, Persson S (2015) A Mechanism for Sustained Cellulose Synthesis during Salt Stress. Cell 162: 1353–1364

Fang L, Ishikawa T, Rennie EA, Murawska GM, Lao J, Yan J, Tsai AY-L, Baidoo EEK, Xu J, Keasling JD, Demura T, Kawai-Yamada M, Scheller HV, Mortimer JC (2016) Loss of Inositol Phosphorylceramide Sphingolipid Mannosylation Induces Plant Immune Responses and Reduces Cellulose Content in Arabidopsis. The Plant Cell 28: 2991–3004

Fenderson BA, Ostrander GK, Hausken Z, Radin NS, Hakomori S-I (1992) A ceramide analogue (PDMP) inhibits glycolipid synthesis in fish embryos. Experimental Cell Research 198: 362–366

Gronnier J, Germain V, Gouguet P, Cacas J-L, Mongrand S (2016) GIPC: Glycosyl Inositol Phospho Ceramides, the major sphingolipids on earth. Plant Signaling & Behavior 11: e1152438

Gu Y, Kaplinsky N, Bringmann M, Cobb A, Carroll A, Sampathkumar A, Baskin TI, Persson S, Somerville CR (2010) Identification of a cellulose synthase-associated protein required for cellulose biosynthesis. Proceedings of the National Academy of Sciences 107: 12866–12871

Gutierrez R, Lindeboom JJ, Paredez AR, Emons AMC, Ehrhardt DW (2009) Arabidopsis cortical microtubules position cellulose synthase delivery to the plasma membrane and interact with cellulose synthase trafficking compartments. Nature Cell Biology 11: 797–806

Inokuchi JI, Momosaki K, Shimeno H, Nagamatsu A, Radin NS (1989) Effects of D-threo- PDMP, an inhibitor of glucosylceramide synthetase, on expression of cell surface glycolipid antigen and binding to adhesive proteins by B16 melanoma cells. Journal of Cellular Physiology 141: 573–583

Ishikawa T, Fang L, Rennie EA, Sechet J, Yan J, Jing B, Moore W, Cahoon EB, Scheller HV, Kawai-Yamada M, Mortimer JC (2018) GLUCOSAMINE INOSITOLPHOSPHORYLCERAMIDE TRANSFERASE1 (GINT1) Is a GlcNAc-Containing Glycosylinositol Phosphorylceramide Glycosyltransferase. Plant Physiology 177: 938–952

Kimberlin AN, Majumder S, Han G, Chen M, Cahoon RE, Stone JM, Dunn TM, Cahoon EB (2013) Arabidopsis 56–Amino Acid Serine Palmitoyltransferase-Interacting Proteins Stimulate Sphingolipid Synthesis, Are Essential, and Affect Mycotoxin Sensitivity. The Plant Cell 25: 4627–4639

Kopytova AE, Rychkov GN, Nikolaev MA, Baydakova GV, Cheblokov AA, Senkevich KA, Bogdanova DA, Bolshakova OI, Miliukhina IV, Bezrukikh VA, Salogub GN, Sarantseva SV, Usenko TC, Zakharova EY, Emelyanov AK, Pchelina SN (2021) Ambroxol increases glucocerebrosidase (GCase) activity and restores GCase translocation in primary patient-derived macrophages in Gaucher disease and Parkinsonism. Parkinsonism & Related Disorders 84: 112–121

Krüger F, Krebs M, Viotti C, Langhans M, Schumacher K, Robinson DG (2013) PDMP induces rapid changes in vacuole morphology in Arabidopsis root cells. Journal of Experimental Botany 64: 529–540

Kyle, Edgar, Jonathan (2016) Substrate specificity, kinetic properties and inhibition by fumonisin B1 of ceramide synthase isoforms from Arabidopsis. Biochemical Journal 473: 593–603

Li S, Ge FR, Xu M, Zhao XY, Huang GQ, Zhou LZ, Wang JG, Kombrink A, McCormick S, Zhang XS, Zhang Y (2013) Arabidopsis COBR-LIKE 10, a GPI-anchored protein, mediates directional growth of pollen tubes. The Plant Journal 74: 486–497

Li S, Lei L, Somerville CR, Gu Y (2012) Cellulose synthase interactive protein 1 (CSI1) links microtubules and cellulose synthase complexes. Proceedings of the National Academy of Sciences 109: 185–190

Liu L, Shang-Guan K, Zhang B, Liu X, Yan M, Zhang L, Shi Y, Zhang M, Qian Q, Li J, Zhou Y (2013) Brittle Culm1, a COBRA-Like Protein, Functions in Cellulose Assembly through Binding Cellulose Microfibrils. PLoS Genetics 9: e1003704

Markham JE, Jaworski JG (2007) Rapid measurement of sphingolipids from Arabidopsis thaliana by reversed-phase high-performance liquid chromatography coupled to electrospray ionization tandem mass spectrometry. Rapid Communications in Mass Spectrometry 21: 1304–1314

Markham JE, Molino D, Gissot L, Bellec Y, Hématy K, Marion J, Belcram K, Palauqui J- C, Satiat-Jeunemaître B, Faure J-D (2011) Sphingolipids Containing Very-Long-Chain Fatty Acids Define a Secretory Pathway for Specific Polar Plasma Membrane Protein Targeting in Arabidopsis. The Plant Cell 23: 2362–2378

Melser S, Batailler B, Peypelut M, Poujol C, Bellec Y, Wattelet-Boyer V, Maneta-Peyret L, Faure J-D, Moreau P (2010) Glucosylceramide Biosynthesis is Involved in Golgi Morphology and Protein Secretion in Plant Cells. Traffic 11: 479–490

Msanne J, Chen M, Luttgeharm KD, Bradley AM, Mays ES, Paper JM, Boyle DL, Cahoon RE, Schrick K, Cahoon EB (2015) Glucosylceramides are critical for cell-type differentiation and organogenesis, but not for cell viability in Arabidopsis. The Plant Journal 84: 188–201

Paredez AR, Persson S, Ehrhardt DW, Somerville CR (2008) Genetic Evidence That Cellulose Synthase Activity Influences Microtubule Cortical Array Organization. Plant Physiology 147: 1723–1734

Paredez AR, Somerville CR, Ehrhardt DW (2006) Visualization of Cellulose Synthase demonstrates functional associate with microtubules. Science 312: 5

Purushotham P, Cho SH, Díaz-Moreno SM, Kumar M, Nixon BT, Bulone V, Zimmer J (2016) A single heterologously expressed plant cellulose synthase isoform is sufficient for cellulose microfibril formation in vitro. Proceedings of the National Academy of Sciences 113: 11360–11365

Purushotham P, Ho R, Zimmer J (2020) Architecture of a catalytically active homotrimeric plant cellulose synthase complex. Science 369: 1089–1094

Rennie EA, Ebert B, Miles GP, Cahoon RE, Christiansen KM, Stonebloom S, Khatab H, Twell D, Petzold CJ, Adams PD, Dupree P, Heazlewood JL, Cahoon EB, Scheller HV (2014) Identification of a Sphingolipid α-Glucuronosyltransferase That Is Essential for Pollen Function in Arabidopsis. The Plant Cell 26: 3314–3325

Roudier FO, Fernandez AG, Fujita M, Himmelspach R, Borner GHH, Schindelman G, Song S, Baskin TI, Dupree P, Wasteneys GO, Benfey PN (2005) COBRA, an Arabidopsis Extracellular Glycosyl-Phosphatidyl Inositol-Anchored Protein, Specifically Controls Highly Anisotropic Expansion through Its Involvement in Cellulose Microfibril Orientation. The Plant Cell 17: 1749–1763

Rugen MD, Vernet MMJL, Hantouti L, Soenens A, Andriotis VME, Rejzek M, Brett P, Van Den Berg RJBHN, Aerts JMFG, Overkleeft HS, Field RA (2018) A chemical genetic screen reveals that iminosugar inhibitors of plant glucosylceramide synthase inhibit root growth in Arabidopsis and cereals. Scientific Reports 8

Smith DK, Jones DM, Lau JBR, Cruz ER, Brown E, Harper JF, Wallace IS (2018) A Putative Protein O-Fucosyltransferase Facilitates Pollen Tube Penetration through the Stigma–Style Interface. Plant Physiology 176: 2804–2818

Stirnemann J, Belmatoug N, Camou F, Serratrice C, Froissart R, Caillaud C, Levade T, Astudillo L, Serratrice J, Brassier A, Rose C, Billette De Villemeur T, Berger M (2017) A Review of Gaucher Disease Pathophysiology, Clinical Presentation and Treatments. International Journal of Molecular Sciences 18: 441

Tartaglio V, Rennie EA, Cahoon R, Wang G, Baidoo E, Mortimer JC, Cahoon EB, Scheller HV (2017) Glycosylation of inositol phosphorylceramide sphingolipids is required for normal growth and reproduction in Arabidopsis. The Plant Journal 89: 278–290

Ternes P, Feussner K, Werner S, Lerche J, Iven T, Heilmann I, Riezman H, Feussner I (2011) Disruption of the ceramide synthase LOH1 causes spontaneous cell death in Arabidopsis thaliana. New Phytologist 192: 841–854

Thevenaz P, Ruttimann UE, Unser M (1998) A pyramid approach to subpixel registration based on intensity. IEEE Transactions on Image Processing 7: 27–41

Updegraff DM (1969) Semimicro determination of cellulose inbiological materials. Analytical Biochemistry 32: 420–424

Vain T, Crowell EF, Timpano H, Biot E, Desprez T, Mansoori N, Trindade LM, Pagant S, Robert S, Höfte H, Gonneau M, Vernhettes S (2014) The Cellulase KORRIGAN Is Part of the Cellulose Synthase Complex. Plant Physiology 165: 1521–1532

Verbančič J, Huang JJ, McFarlane HE (2021) Analysis of cellulose synthase activity in Arabidopsis using spinning disk microscopy. STAR Protocols 2: 100863

Villalobos JA, Yi BR, Wallace IS (2015) 2-Fluoro-L-Fucose Is a Metabolically Incorporated Inhibitor of Plant Cell Wall Polysaccharide Fucosylation. PLOS ONE 10: e0139091

Wang W, Yang X, Tangchaiburana S, Ndeh R, Markham JE, Tsegaye Y, Dunn TM, Wang G-L, Bellizzi M, Parsons JF, Morrissey D, Bravo JE, Lynch DV, Xiao S (2008) An Inositolphosphorylceramide Synthase Is Involved in Regulation of Plant Programmed Cell Death Associated with Defense in Arabidopsis. The Plant Cell 20: 3163–3179

Xia Y, Lei L, Brabham C, Stork J, Strickland J, Ladak A, Gu Y, Wallace I, Debolt S (2014) Acetobixan, an Inhibitor of Cellulose Synthesis Identified by Microbial Bioprospecting. PLoS ONE 9: e95245

